# Proteomic study of differentially expressed proteins in seeds between parents and offspring of castor bean (*Ricinus communis* L.)

**DOI:** 10.1101/2022.04.13.488138

**Authors:** Xiaotian Liang, Rui Luo, Yanxin Zhang, Mingda Yin, Yanpeng Wen, Xuemei Hu, Zhiyan Wang, Yumiao Huo, Fenglan Huang

## Abstract

Castor bean (*Ricinus communis L*.), one of the top 10 oilseed crops in the world, has high economic value. Hybridization is the most direct and effective method to breed new varieties with high yield, high oil content, and strong stress resistance. Therefore, prediction of desired traits in castor hybrid offspring is particularly important. In this study, proteomic analysis was performed to identify differentially expressed proteins (DEPs) in seeds between castor hybrid offspring and their female (Lm female line aLmAB2) and male parents (CSR•181). Among the DEPs upregulated in the seeds of hybrid offspring, the majority were related to seed yield and stress resistance, while some were related to oil synthesis and fatty acid synthesis and metabolism in seeds. In other words, the hybrid offspring showed heterosis for seed yield, stress resistance, oil synthesis, and fatty acid synthesis and metabolism when compared with their parents. Further, real-time quantitative polymerase chain reaction assays were performed on 12 genes encoding DEPs involved in oil synthesis, pollen abortion, yield, and stress resistance of seeds. The results showed that the expression levels of the 12 genes were consistent with those of the DEPs.

## Introduction

Castor bean (*Ricinus communis* L.) belongs to the genus *Ricinus* in the family Euphorbiaceae. Due to its strong environmental adaptability, castor bean can grow in harsh conditions such as saline-alkali soil and heavy metal–contaminated soil and is considered a valuable plant resource for phytoremediation. Therefore, it has been widely planted in tropical, subtropical, and warm temperate countries, especially in India, China, and Brazil [1,2]. China has more than 2300 germplasm resources of castor collected both at home and abroad, and in-depth studies have been conducted to further explore the application of castor bean. In Inner Mongolia, castor bean is a characteristic oilseed crop pivotal to local economic and social development [3]. It is a special industrial oil crop with prominent intraspecific heterosis.

Seed storage substances gradually accumulate during seed development and maturation. Protein, one of the seed storage substances, can help a seed to form cellular structures and provide a nitrogen source for normal metabolism and development of seeds, thereby playing a vital role in cellular structure and function [4]. In 1994, Wilkins and Williams first proposed the concept of the proteome, which refers to all proteins expressed in a cell, tissue, or organism [5]. The methods used for proteomic analysis of seeds mainly include two categories: gel and nongel analysis. Gel analysis is most commonly used, including two-dimensional gel electrophoresis (2-DE) and two-dimensional difference gel electrophoresis (2D-DIGE) [6]. After the differentially expressed spots are determined by using 2-DE and 2D-DIGE, a complete protein atlas can be obtained by mass spectrometry (MS), and this can better reflect the changes in protein expression and posttranslational modification [7,8]. Proteomic studies of plant seeds have attracted an increasing amount of attention because they can provide information on substance changes in seeds during their growth and development and help explore the adaptative mechanism of plants to environmental stress. Xia et al. [9] conducted proteomic analysis to investigate the impact of temperature on seed metabolism in dormant and germinating seeds and deciphered the regulation of central metabolism activity during seed germination. Gu et al. [10] studied seed germination vigor in *Brassica napus* seeds differing in oil content by an integrated proteomic and genomic approach, and their research laid a foundation for studying the regulatory mechanism of seed germination vigor in *Brassica napus*.

In this study, the seeds from castor female parents (the Lm female line aLmAB2), male parents (CSR•181), and their hybrid offspring were used as the study materials, and the heterotic traits of offspring compared with their parents were identified by proteomic techniques. Further, genes associated with these traits were determined and tested with reverse transcription–quantitative PCR (RT-qPCR). The findings of this study can lay a foundation for guiding the production practice of castor bean and cultivating new castor varieties with high yield, high oil content, and strong stress resistance.

## Materials and methods

### Experimental materials

#### Plant materials

The plant materials used in this study were the castor seeds from female parent (aLmAB2), male parent (CSR•181), and their hybrid offspring F_0_ and F_1_. Castor plants at the flowering stage were bagged and pollinated manually. Subsequently, the seeds that had a developmental time of 20 days (Fig 1A), 40 days (Fig 1B), and 60 days post pollination (DPP) (Fig 1C) were collected. After their testa were removed, the seeds were wrapped in tinfoil paper and stored in an ultralow-temperature freezer at −80°C for subsequent use. The experimental materials were provided by Tongliao Research Institute of Agriculture and Animal Husbandry.

**Fig 1.**
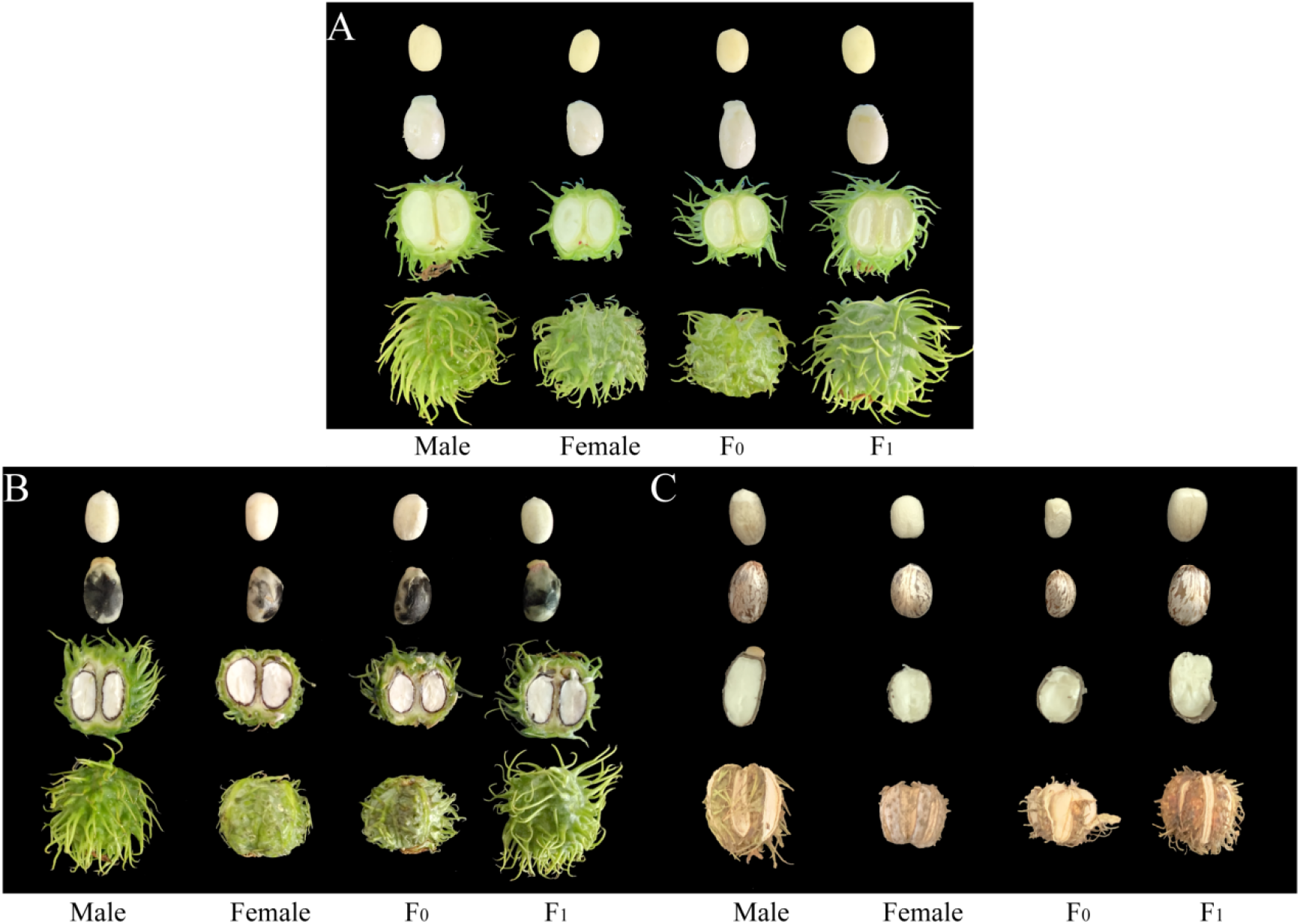
Castor Seeds at Different Developmental Stages. Note: A: 20 DPP; B: 40 DPP; C: 60 DPP

#### Reagents and instruments

The reagents used in this experiment included Tris-HCI solution, ethylene diamine tetraacetic acid stock solution, 0.4% β-mercaptoethanol solution, 0.9 M sucrose, balanced phenol solution, extraction solution, ammonium acetate/methanol solution, acetone solution, lysis solution, NaOH solution, HCl solution, lysine solution, Amersham CyDye DIGE Fluors (minimal dyes) for the Ettan DIGE kit (GE), the 2-D Clean-up kit (GE), the 2-D Quant kit (GE), hydration solution, 87% glycerol, covering oil, monomer storage solution, 4× separating gel buffer, 10% sodium dodecyl sulfate (SDS) solution, N,N,N′,N′-tetramethylethylenediamine (TEMED), 10% ammonium persulfate solution, 75% alcohol, gel-sealing liquid, equilibrium solution, bromophenol blue dye, 1,4-dithiothreitol (DTT), iodoacetamide (IAA), premixed protein marker, SDS electrophoresis buffer, stationary liquid, amplification solution, Coomassie brilliant blue dye, decolorizing solution, RNA Rapid Extraction Kit for plants (ZOMANBIO), GoTaq® qPCR and RT-qPCR Systems (Promega), and reverse transcription system (Promega). All reagent formulas are shown in S1 Table. The instruments used in this study included high-speed and low-speed refrigerated centrifuges (Thermo), an Infinite M200 Pro multifunctional microplate reader (TECAN), a gel imaging system and electrophoresis apparatus (Bio-Rad), the Ettan IPGphor 3 for electrophoresis by isoelectric focusing (IEF) in the first dimension, a MultiTemp IV water bath, a FLA9500 multifunctional biomolecular imager (GE), and a real-time fluorescence-based quantitative PCR system (ABI 7500, ABI company).

### 2-DE

#### Sample preparation

An optimized phenol extraction method was used to extract protein from castor seeds. The obtained dry protein powder was aliquoted into 1.5-mL centrifuge tubes. Each tube was added with a 20-fold volume of lysis solution (each 1 mL of lysis solution was added to 0.0062 g DTT, 20 μL of immobilized pH gradient (IPG) buffer, and 40 μL of protease inhibitor before use). Then, the tube was vortexed at a low temperature for 30 minutes, followed by an ultrasound ice-bath for 20 minutes. The process was repeated until dry protein powder was fully dissolved. After centrifugation at 13,000*g* for 1 hour at 4 °C, the supernatant was transferred to a 1.5-mL centrifuge tube. The 2-D Clean-up kit was used to remove impurity from the sample. The 2-D Quant kit was used for sample quantitation, which derived a standard curve: y = −0.0041x + 0.3751 (R^2^ = 0.999), where y was the absorbance value of the postquantitation sample at 480 nm and x was the protein concentration (μg/μL).

#### 2-DE procedures

For the first dimension, 450 μL of the sample solution containing 800 μg of protein was loaded for electrophoresis by IEF. First, 800 μg of the lysed sample was transferred into a centrifuge tube, which was then added to hydration solution to achieve a total volume of 450 μL (each 1 mL of hydration solution was added with 5 μL of NL3-10 IPG buffer and 0.0028 g of DTT). Then, the sample solution was vacuum-centrifuged at 13,000 rpm for 1 hour at 4 °C. After the centrifuged sample solution was transferred to the strip groove, a 24-cm-long dry gel strip (pH 3-10 NL) was placed into the groove; after ensuring that the gel surface was fully in contact with the sample, the covering oil was added, and then a lid was placed to cover the gel strip groove. Finally, after all gel strips of samples had been placed into a PROTEAN IEF instrument, the following IEF protocol was run the Table 1 at a constant temperature of 20 °C:

**Table 1.** IEF Program.

| Program | Voltage/V | Time/h | Purpose |
| --- | --- | --- | --- |
| S1 | 30 | 8 | Hydration |
| S2 | 50 | 4 | Desalination |
| S3 | 100 | 1 | Desalination |
| S4 | 300 | 1 | Desalination |
| S5 | 600 | 1 | Desalination |
| S6 | 1000 | 2 | Desalination |
| S7 | 8000 | 12 | Focusing |
| S8 | 1000 | ∞ | Maintaining |

For the second-dimension SDS-PAGE, 12.5%T polyacrylamide gel was used. First, the gel strips after the first-dimension IEF were equilibrated for 15 minutes in each of 2% (w/v) DTT and 2.5% (w/v) IAA equilibration buffer. Then the well-equilibrated strips were sealed with sealing liquid in a gel plate, which was then placed in the Ettan DALT 12 unit (12 gels per run) (GE-061-TY7119, GE Healthcare). Electrophoresis was performed at 2 W per gel for 1 hour and then 16 W per gel until the bromophenol blue band was 0.5-1 cm away from the bottom of the gel.

After electrophoresis, the gel was taken out and sequentially submerged in stationary liquid for 30 minutes, amplification solution for 20 minutes, Coomassie brilliant blue dye for 10 minutes, and finally the decolorizing solution until only protein spots appeared on the gel, without other, interfering dye residues. The Image Scanner III image scanning system (28-9076-07 GE Healthcare, USA) was used for gel imaging, and the Image Master 2-D Platinum 7.0 software was used for gel image analysis.

### 2D-DIGE

#### Sample preparation

The sample was lysed using the lysis solution without DTT or IPG buffer. After lysis, the sample solution was adjusted to pH 8.0-9.0 using 100 mM NaOH and 1 M HCl, with a target pH of 8.5. The subsequent procedures were the same as those described in 1.2.1.

#### Preparation of CyDye DIGE fluor minimal dyes

Preparation of 1 nmol/μL fluorescent dye stock solution: After the dye-storing tube was taken out of the −80°C refrigerator, each tube received 5 μL of *N,N*-dimethylformamide solution and then vortexed for 30 seconds to fully dissolve the dye. Next, it was centrifuged at 12,000*g* for 30 seconds to push the dye stock solution to the bottom of the tube. Preparation of 400 pmol/μL fluorescent dye working solution: Five microliters of fluorescent dye stock solution was diluted with 7.5 μL of *N,N*-dimethylformamide solution to achieve a final concentration of 400 pmol/μL, followed by centrifugation at 12,000*g* for 30 seconds to push the solution to the bottom of the tube. All the above procedures were performed at 4 °C in the dark.

A certain volume of the sample solution (containing 50 μg of protein) was taken out and mixed with 1 μL of the dye working solution through vortexing, followed by a brief centrifugation. Then, as a quench-labeling reaction, the sample was kept on ice in the dark for 30 minutes and then mixed with 1 μL of 10 mM lysine solution through vortexing. After a brief centrifugation, the sample was placed on ice in the dark for 10 minutes.

#### 2D-DIGE procedures

According to the experimental design, the internal standard sample and the target samples labeled with Cy2, Cy3, or Cy5 were mixed in a centrifuge tube and then added to hydration solution (every 1 mL of hydration solution had been added with 5 μL of NL3-10 IPG buffer and 0.0028 g of DTT) up to a total volume of 450 μL. After being placed on ice in the dark for 10 minutes, the sample solution was treated as in 1.2.2 for the first-dimension IEF.

For the second-dimension SDS-PAGE, 12.5% polyacrylamide gel was prepared using low-fluorescence glass plates, followed by the same procedures as in 1.2.2. Light exposure was avoided during the entire process. The Typhoon FLA9500 multifunctional biomolecular imager was used for gel imaging, and the DeCyder software for 2D-DIGE analysis was used to identify and annotate the differentially expressed spots (ratio ≥ 1.5 and ANOVA ≤ 0.05).

### MS identification and analysis of the DEP spots

#### MS identification

The DEP spots in the gel were recovered and sent to Shanghai Bioclouds Biological Tech Co. Ltd. for MS identification.

#### Database search and functional classification of differentially expressed proteins (DEPs)

The information of the DEPs identified by MS were obtained by searching the database of castor bean proteins (https://www.ncbi.nlm.nih.gov/). The proteins were functionally classified into Gene Ontology (GO) categories according to UniProt (http://www.UniProt.org/) and QuickGO (https://www.ebi.ac.uk/QuickGO/), and their roles in metabolic pathways were categorized according to the Kyoto Encyclopedia of Genes and Genomes (KEGG) Pathway Database (https://www.kegg.jp/kegg/pathway.html). The histograms of GO classification were generated on the Omics Cloud platform established by GENE DENOVO (http://www.omicshare.com/index.php). The heatmaps were drawn using GraphPad Prism7.04 software.

### RT-qPCR verification

The RNA Rapid Extraction Kit for plants was used to extract RNA from castor seeds. The extracted RNA was reverse-transcribed into cDNA. SnapGene 2.8.0 software was used to design primers (Table 2).

**Table 2.**
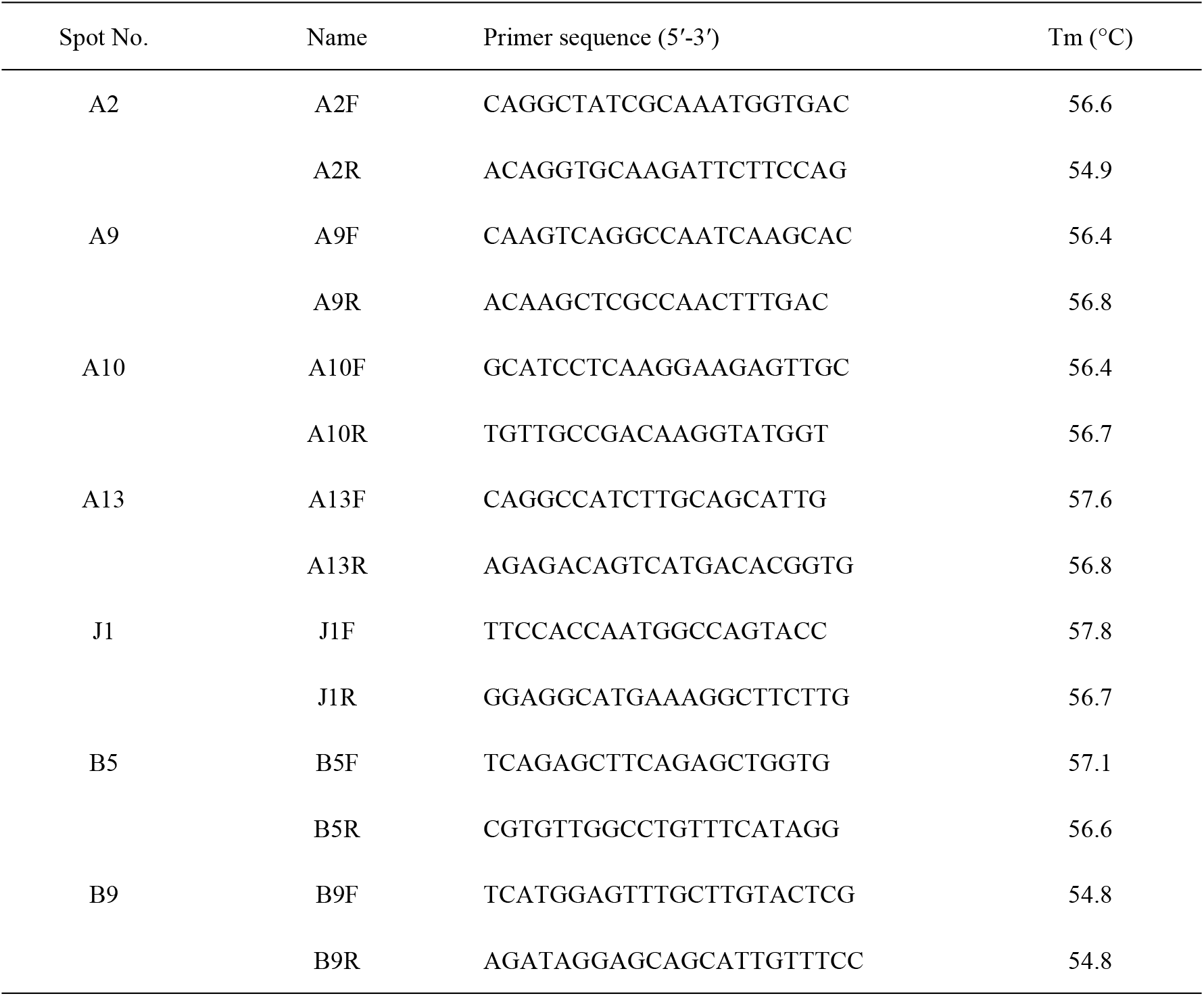
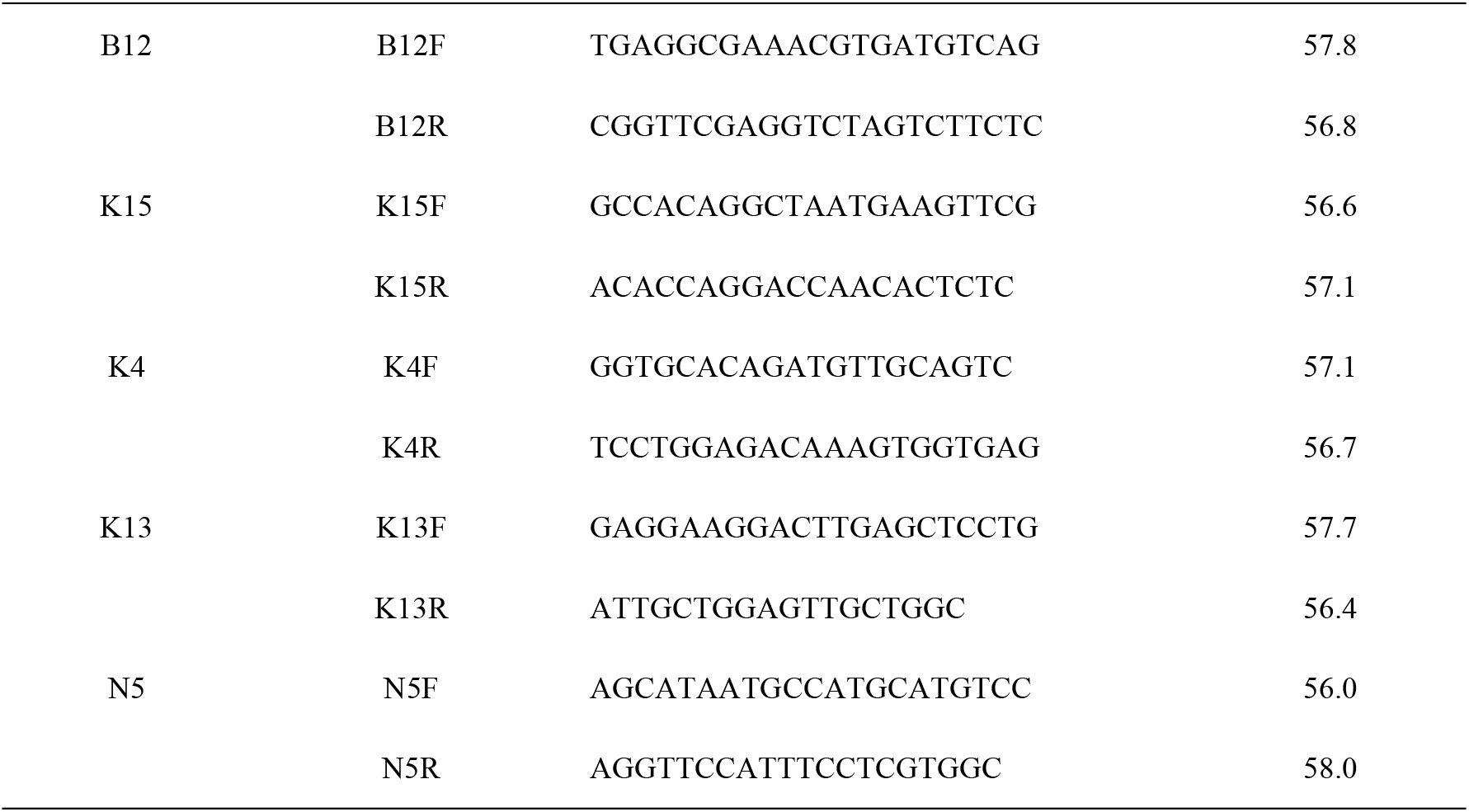
RT-qPCR Primer Sequences.

The GoTaq® qPCR systems was used for qPCR. The reaction system consisted of 10 μL of SYBR Premix Ex *Taq*, 0.4 μL of Rox reference dye II, 1.0 μL of each of the upstream and downstream primers, 2.0 μL of template DNA, and ddH_2_O up to 20 µL. The amplification program was by 45 cycles of 95°C for 30 seconds (s), 95°C 10 s, melting temperature for 45 s, and 72 °C for 10 s. The results were calculated using the 2-^ΔΔCt^ method.

One-way ANOVA was performed with SPSS version 19.0 software, and the relevant histograms were generated in Excel 2019 software.

## Results

### Optimization of the 2-DE system

#### Optimization of conditions for 2-DE

To determine whether it is necessary to remove impurities from protein samples in this experiment, the same protein samples were divided into two groups, one with impurity removal and the other without impurity removal, before 2-DE (24-cm dry gel strips, pH 3-10 NL, protein loading amount 800 μg). The results are presented in S1 Fig. To determine the optimal pH range for this experiment, the protein samples with the same treatment (with or without impurity removal) were subjected to IEF and SDS-PAGE using the 24-cm dry gel strips with pH 3-10 NL or pH 4-7. The results are shown in S2 Fig. To determine the optimal protein loading amount, different amounts of protein (800 μg, 1000 μg, and 1200 μg) were loaded for 2-DE (24-cm dry gel strips; pH 3-10 NL) under the same conditions. The comparison results are shown in S3 Fig. Based on the results of these preliminary experiments, the optimal experimental conditions were impurity removal, 24-cm dry gel strips with pH 3-10 NL, and a loading amount of 800 μg. These conditions were used in the following experiments.

### 2D-DIGE results

The castor seeds at different developmental stages from parents and offspring were subjected to 2D-DIGE. The results are shown in Fig 2.

**Fig 2.**
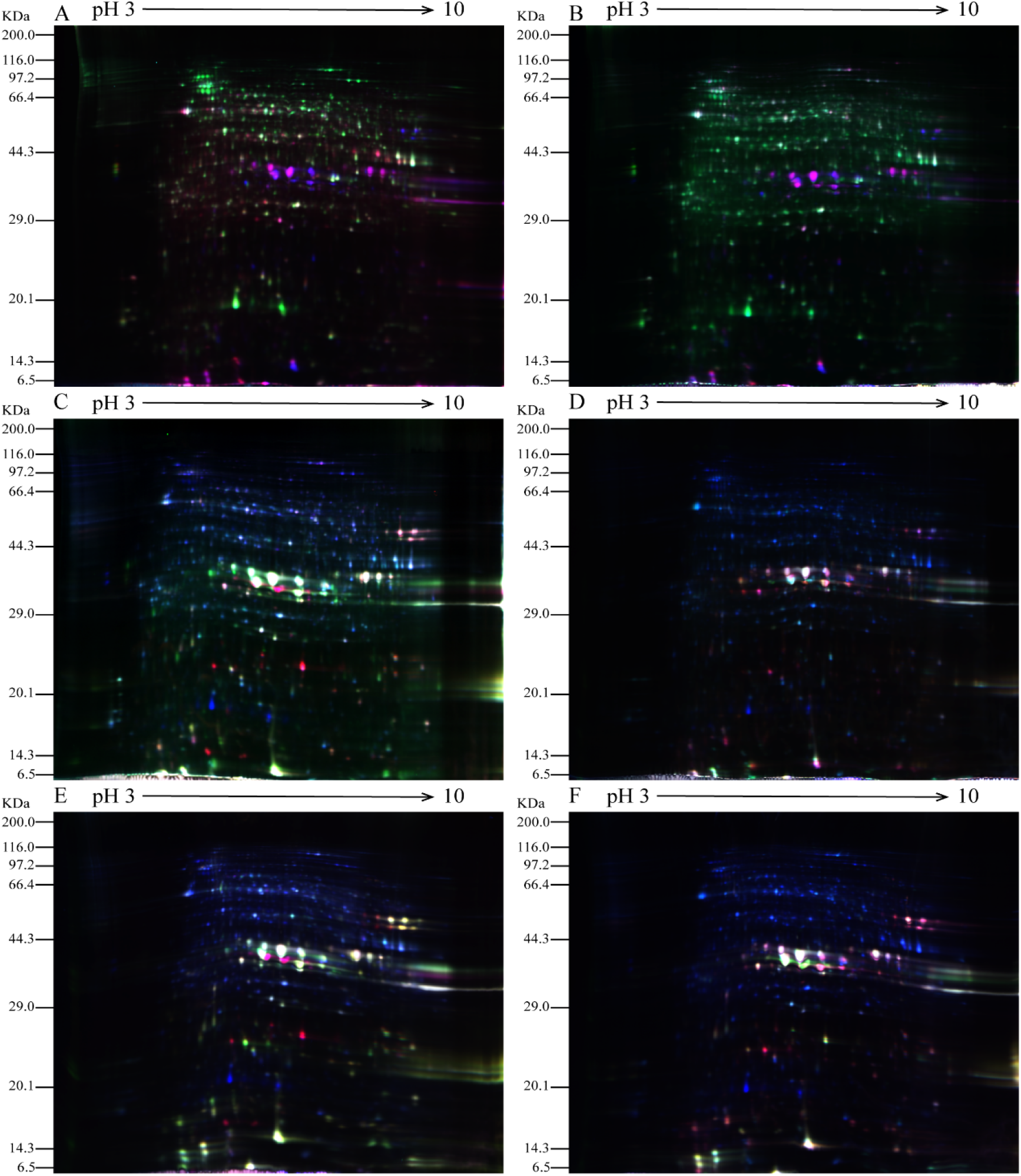
2D-DIGE Results of Castor Parents and their Offspring. Note: A: Male, F_0_, 20 d. B: Female, F_1_, 20 d, A&B when they had Developed to 20 DPP. C: Male, F_0_, 40 d. D: Female, F_1_, 40 d, C&D when they had Developed to 40 DPP. E: Male, F_0_, 60 d. F: Female, F_1_, 60 d, E&F when they had Developed to 40 DPP.

In the figures, the blue protein spots denote the internal standard samples labeled with Cy2; the green protein spots are the Cy3-labeled samples from male and female parents; and the red protein spots are the Cy5-labeled samples from castor offspring. In Fig 2, the 2D-DIGE protein spots for the seeds at different development stages from both castor parents and offspring indicate that the dye labeling of all three samples was successful and met the requirements for experimental analysis.

### MS identification and functional classification results of DEPs

#### MS identification results

The 2D-DIGE results of castor seeds at different development stages from parents and their offspring were analyzed for differences using DeCyder 2-D software. A total of 122 differentially expressed spots (ratio ≥ 1.5 and ANOVA ≤ 0.05) were determined and recovered, of which 121 spots containing 93 proteins were successfully identified by matrix-assisted laser desorption/ionization mass spectrometry (MALDI-TOF-MS). The results are shown in S2 Table. The sources/names and classification of these DEPs are listed in S3 Table. Among the DEPs between the castor parents and their offspring, 69 were present in seeds at 20 DPP, 46 in seeds at 40 DPP, and 20 in seeds at 60 DPP. Some proteins had a significantly different expression in the castor seeds at two or all three development stages between castor parents and their offspring, such as starch synthase, D-7 late embryogenesis abundant (LEA) protein, superoxide dismutase (SOD), and aspartic proteases (APs).

#### Functional classification results of the DEPs

The 93 DEPs identified were classified to five categories (Fig 3) related to stress responses (61.29%), seed nutrient storage (17.20%), fatty acid synthesis and metabolism (12.90%), amino acid metabolism (6.45%), and others (2.15%). The DEPs were most often related to stress responses. These included SOD, APs, D-7 LEA protein, and glutathione peroxidase. The DEPs associated with nutrient storage in seeds included starch synthase (chloroplast/starch), 2S albumin precursor, and vicilin GC72-A. The DEPs associated with fatty acid synthesis and metabolism included stearoyl-ACP desaturase, oil body–associated protein 2C (OBAP2C), and acetyl-CoA carboxylase. The DEPs associated with amino acid metabolism included transaldolase, transketolase (chloroplast), S-adenosylmethionine synthase (SAMS), and others.

**Fig 3.**
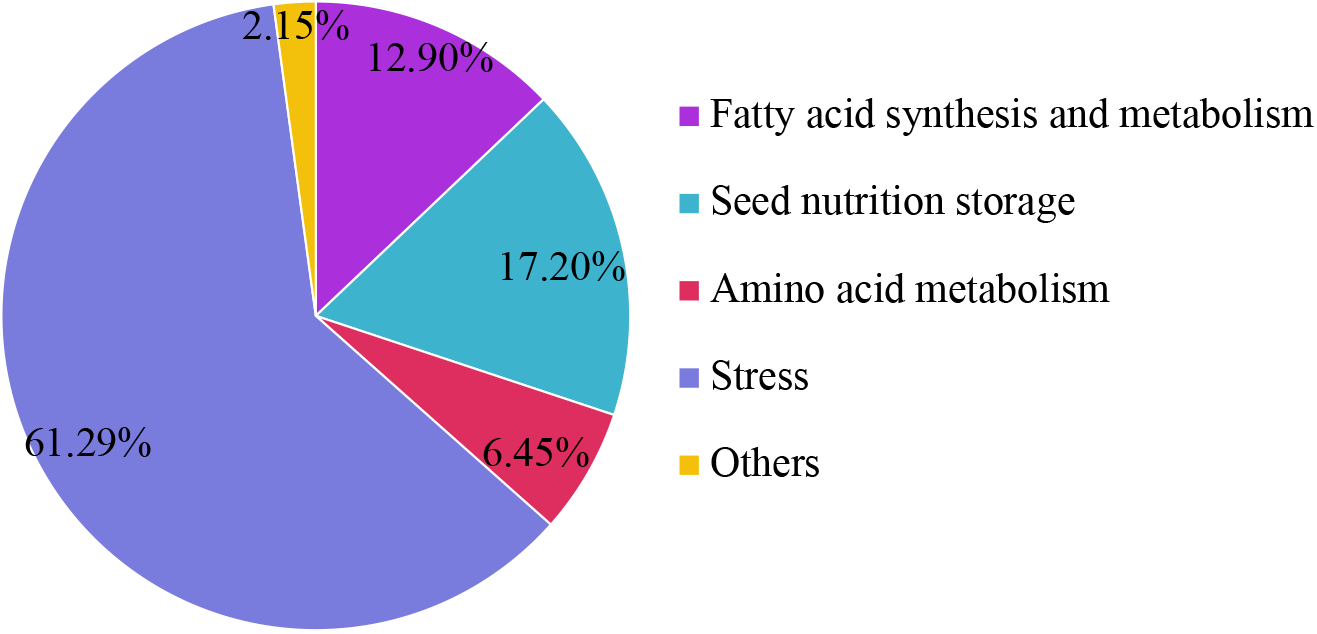
Functional Classification of the Differentially Expressed Proteins between Castor Parents and their Offspring.

##### Functional classification results of the DEPs in seeds at 20 DPP between castor parents and their offspring

A total of 69 DEPs were identified in seeds at 20 DPP between castor parents and their offspring. The seeds from F_1_ offspring had 16 upregulated DEPs (e.g., OBAP2C, acetyl transferase NATA1, and 2S albumin precursor) when compared with their male parent and 13 upregulated DEPs (e.g., protein disulfide isomerase (PDI), luminal-binding protein 5, and triphosphate isomerase) when compared with their female parent. The seeds from F_1_ offspring had 23 downregulated DEPs (e.g., dihydrolipoyl dehydrogenase 1, MLP-like protein 34, and salicylic acid-binding protein 2) when compared with their male parent and nine downregulated DEPs (e.g., l-ascorbate peroxidase, adenosine kinase 2, and the 36-kDa porin in mitochondrial outer membrane) when compared with their female parent. In the F_0_ seeds, 11 DEPs (e.g., legume protein A, SOD, and hydroxyl-ACP dehydrase) were upregulated when compared with their male parent, and five DEPs (i.e., transaldolase, stearoyl-ACP desaturase, 60S acidic ribosomal protein P2, SOD, and uncharacterized protein) were upregulated when compared with their female parent. The F_0_ seeds had 11 downregulated DEPs (e.g., the major latex allergen Hev b5, MLP-like protein 31, and GDP-mannose 3,5-epimerase 1) when compared with their male parent and four downregulated DEPs (i.e., adenosyl homocysteinase, xylem serine proteinase 1 precursor, cytochrome *c* oxidase subunit 6b-1, and lactoylglutathione lyase GLX1) when compared with their female parent. The relative expression levels of DEPs are shown in S4 Fig, and the results regarding their involvement in KEGG metabolic pathways are shown in S5 Fig.

Fig 4 shows the GO functional annotations of the 69 DEPs in the three categories of biological process, cellular component, and molecular function. The DEPs between F_1_ and parents and DEPs between F_0_ and parents were mostly associated with biological processes, including starch synthase, ribulose bisphosphate carboxylase small chain, and glutathione peroxidase. A smaller number of DEPs were associated with molecular functions, including glutathione peroxidase, APs, and acetyltransferase NATA1. The fewest DEPs were associated with cellular components, including 60S acidic ribosomal protein P2, stearoyl-ACP desaturase, and PDI. The analysis of KEGG metabolic pathways showed that the DEPs between F_1_ and parents and the DEPs between F_0_ and parents were mostly involved in rcu01100 (metabolic pathways) and rcu01110 (biosynthesis of secondary metabolites), such as alcohol dehydrogenase 1 (ADH1), starch synthase, and hydroxyl-ACP dehydrase. A number of DEPs were involved in rcu01230 (biosynthesis of amino acids), rcu00710 (carbon fixation in photosynthetic organisms), and rcu00061(fatty acid biosynthesis).

**Fig 4.**
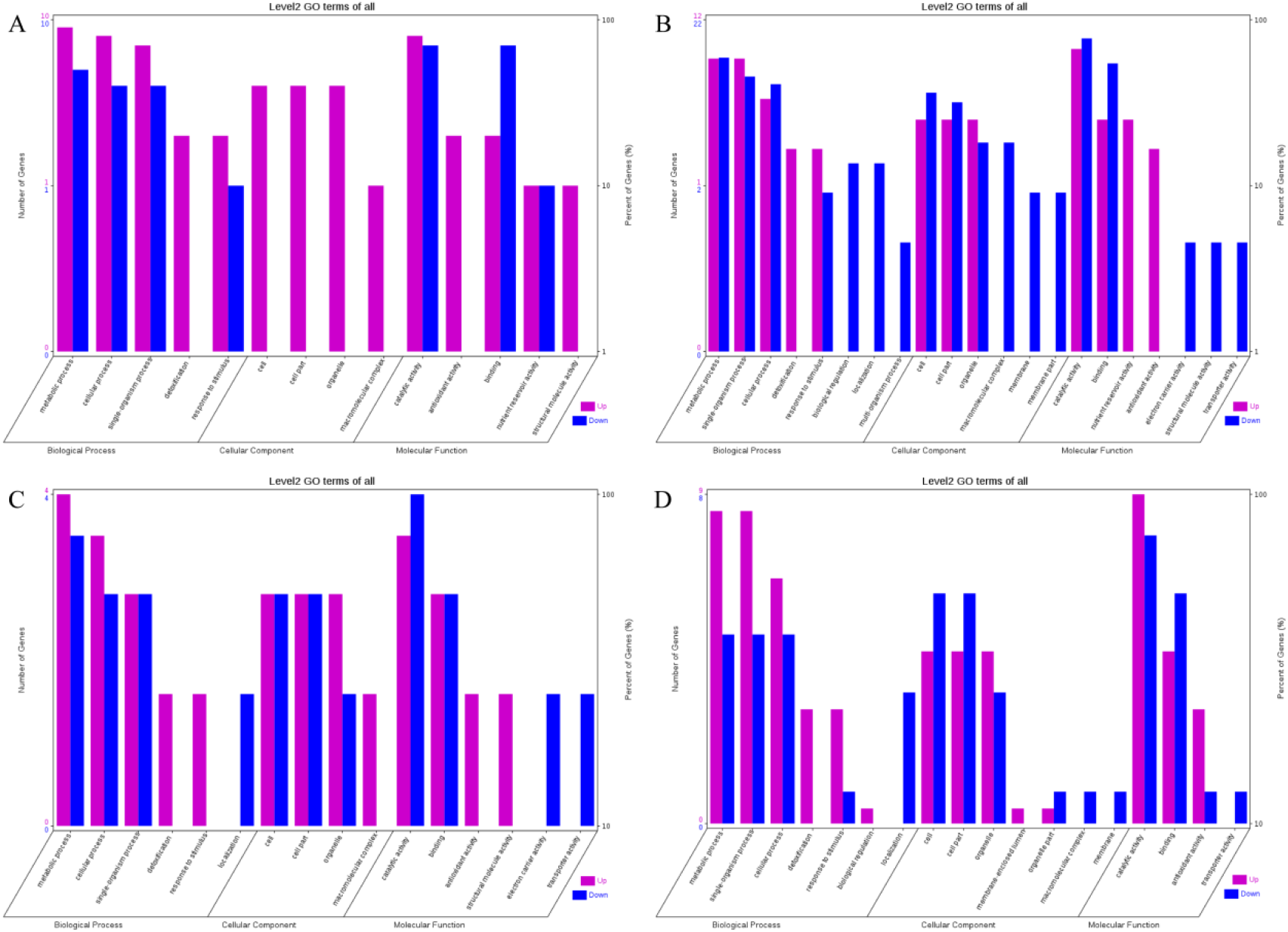
GO Functional Annotation of Differentially Expressed Proteins in Seeds at 20 DPP between Castor Parents and their Offspring. Note: A: Male, F_1_, 20 DPP; B: Female, F_1_, 20 DPP, C: Male, F_0_, 20 DPP; D: Female, F_0_, 20 DPP.

##### Functional classification results of the DEPs in seeds at 40 DPP between castor parents and their offspring

A total of 46 DEPs were identified in seeds at 40 DPP between castor parents and their offspring. The F_1_ seeds had three upregulated DEPs when compared with their male parent, including D-7 LEA protein, legume protein A, and heat shock protein (HSP); and they had 10 upregulated DEPs when compared with their female parent, including the large subunit of 2S sulfur-rich seed storage protein, nucleolar protein 56 (NOP56), and reduced nicotinamide adenine dinucleotide phosphate (NADPH)-dependent aldehyde reductase 1. Compared with their male parents, four DEPs were downregulated in the F_1_ seeds, including ATP synthase subunit β, HSP, ADH1, and plastid 3-keto-acyl-ACP synthase; when compared with female parents, six DEPs were downregulated, including starch synthase, fructose-bisphosphate aldolase, GroES chaperonin, etc. In the F_0_ seeds, seven DEPs (e.g., APs, large subunits of 2S sulfur-rich seed storage protein, and nucleoside diphosphate kinase (NDPK)) and nine DEPs (e.g., SOD, D-7 LEA protein, and glutelin type-A precursor) were upregulated when compared with the male and female parents, respectively. One DEP (enoyl-ACP reductase (EAR) precursor) and three DEPs (i.e., translation-controlled tumor protein homolog, pre-pro-ricin, and GroES chaperonin) were downregulated when compared with the male and female parents, respectively. Compared with the F_0_ generation, the seeds of the F_1_ generation had seven upregulated DEPs (e.g., legume protein A, HSP, and 14-3-3-like protein D isoform X1) and seven downregulated DEPs (e.g., fructose-bisphosphate aldolase 1, adenosine kinase 2, and adenosyl homocysteinase). The relative expression levels of the DEPs mentioned above are depicted in S6 Fig, and the results regarding their involvement in the KEGG metabolic pathways are presented in S7 Fig.

The GO functional annotations for the 46 DEPs are presented in Fig 5. The majority of DEPs in seeds between F_1_ and male parent were related to biological processes and molecular functions; compared with the male parent, only legume protein A was upregulated while all other DEPs were downregulated in the F_1_ seeds. Most of the DEPs in seeds between F_1_ and the female parent (e.g., NADPH-dependent aldehyde reductase 1 and rRNA N-glycosidase) were associated with biological processes and molecular functions. Similarly, most of the DEPs between F_0_ and parents were related to biological processes and molecular functions, including cysteine protease inhibitor and glutelin type-A precursor. The DEPs between F_1_ and F_0_ were mostly related to biological processes, including SAMS and adenosyl homocysteinase; the DEP categorized under cellular component was enolase, and those categorized under molecular function included HSP and 14-3-3-like protein D isoform X1. According to the KEGG metabolic pathways, some HSP family members involved in rcu04141 (protein processing in endoplasmic reticulum), including HSP, were differentially expressed in seeds between F_1_ and parents, between F_0_ and parents, and between F_1_ and F_0_. In addition, among the upregulated DEPs in the F_0_ seeds vs. the male parent, NDPK was involved in rcu04016 (MAPK signaling pathway) and betaine aldehyde dehydrogenase 1 (BADH1) was involved in rcu00260 (glycine, serine, and threonine metabolism). SOD, an upregulated DEP in the F_0_ seeds compared with their female parent, was involved in rcu04146 (peroxisome); and chaperonin, an upregulated DEP in the F_1_ generation compared with F_0_, was involved in rcu03018 (RNA degradation).

**Fig 5.**
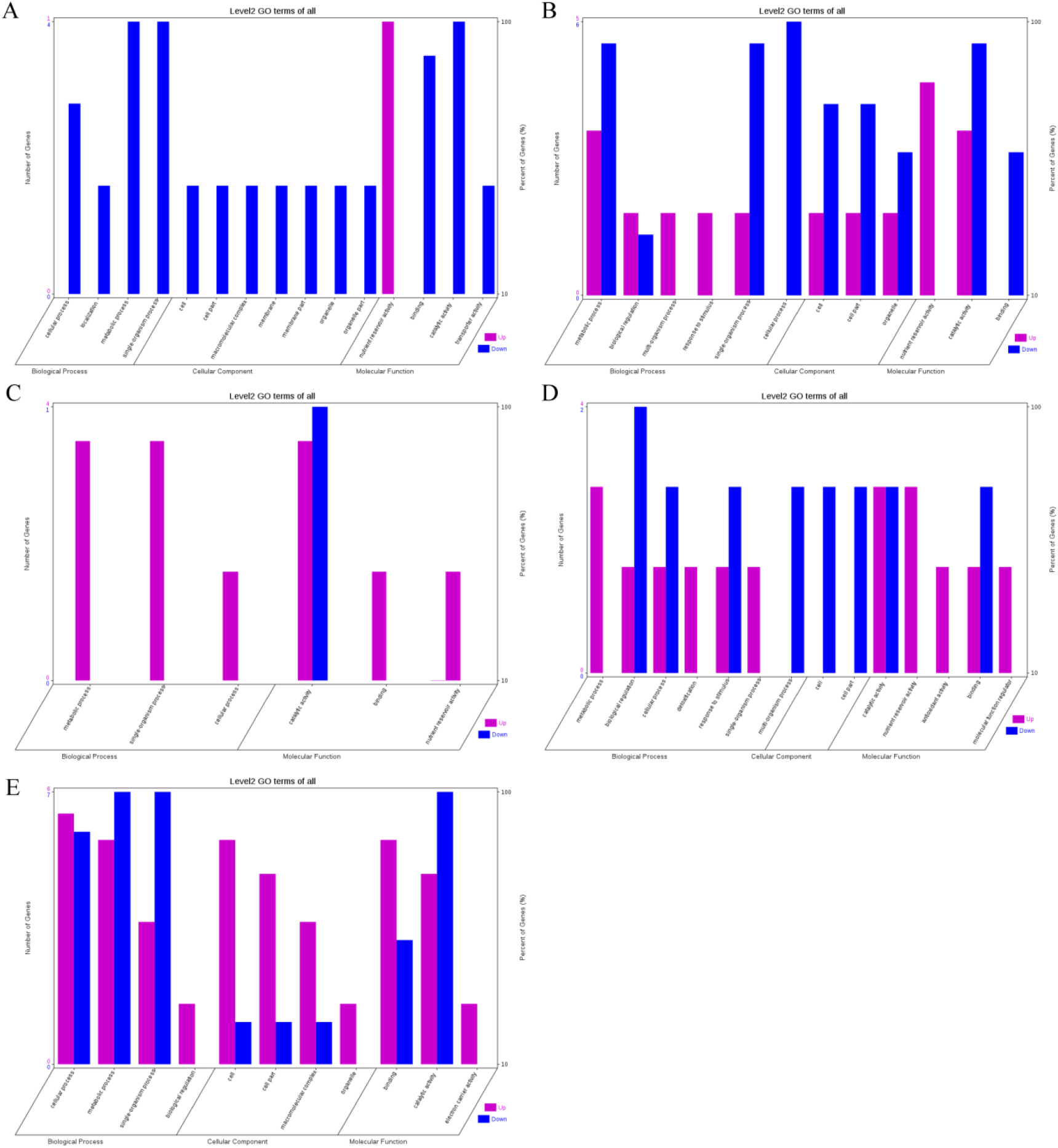
GO Functional Annotation of Differentially Expressed Proteins in Seeds at 40 DPP between Castor Parents and their Offspring. Note: A: Male, F_1_, 40 DPP; B: Female, F_1_, 40 DPP; C: Male, F_0_, 40 DPP; D: Female, F_0_, 40 DPP; E: F_1_ vs. F_0_, 40 DPP.

##### Functional classification results of the DEPs in seeds at 60 DPP between castor parents and their offspring

A total of 25 DEPs were identified in seeds at 60 DPP between castor parents and their offspring. The F_1_ seeds had two upregulated DEPs when compared with their male parent, NOP56 and 5-methyltetrahydroproylglutamate-homocysteine methyltransferase; and they had six upregulated DEPs when compared with their female parent, including D-7 LEA protein, NDPK, and large subunits of ribulose-1,5-bisphosphate carboxylase/oxygenase. Five DEPs were downregulated in the F_1_ seeds compared with their male parent, including malate dehydrogenase, phospho-glucosidase, and enolase, and no DEP showed detectable downregulation in expression in the F_1_ seeds compared to their female parent. In the seeds of the F_0_ generation, three DEPs (i.e., vicilin GC72-A, legume protein A, and seed storage protein) and eight DEPs (e.g., 6-phosphogluconate dehydrogenase, large subunits of 2S sulfur-rich seed storage protein, and D-7 LEA protein) were upregulated compared to the male and female parents, respectively. No DEP showed detectable downregulation in the F_0_ seeds compared with either of their parents. Compared with F_0_, the F_1_ seeds had five upregulated DEPs (e.g., 2S albumin precursor, phospho-glucosidase, and malate dehydrogenase) but no downregulated DEP. The relative expression levels of the abovementioned DEPs are shown in S8 Fig, and the results on their involvement in KEGG metabolic pathways are shown in S9 Fig. The DEPs between F_0_ and the male parent were not involved in any metabolic pathways.

The GO functional annotations of the 25 DEPs are shown in Fig 6. Most of the DEPs between F_1_ and parents and between F_0_ and parents (e.g., ADH1, NOP56, and NDPK) were related to biological processes and molecular functions. Most of the DEPs between F_1_ and F_0_ (e.g., phospho-glucosidase, malate dehydrogenase, and enolase) were related to biological processes. From the KEGG metabolic pathway analysis, the DEPs between F_1_ and parents, between F_0_ and parents, and between F_1_ and F_0_ were mostly involved in rcu01100 (metabolic pathways) and rcu01110 (biosynthesis of secondary metabolites), including 6-phosphogluconate dehydrogenase, ADH1, and 5-methyltetrahydroproylglutamate-homocysteine methyltransferase. In addition, the DEPs upregulated in F_1_ and F_0_ over their parents were also involved in the pathways of rcu00270 (cysteine and methionine metabolism), rcu00592 (alpha-Linolenic acid metabolism), rcu00230 (purine metabolism), etc. The DEPs upregulated in the F1 generation compared with F0 also participated in the pathways related to rcu01200 (carbon metabolism) and rcu01230 (biosynthesis of amino acids).

**Fig 7.**
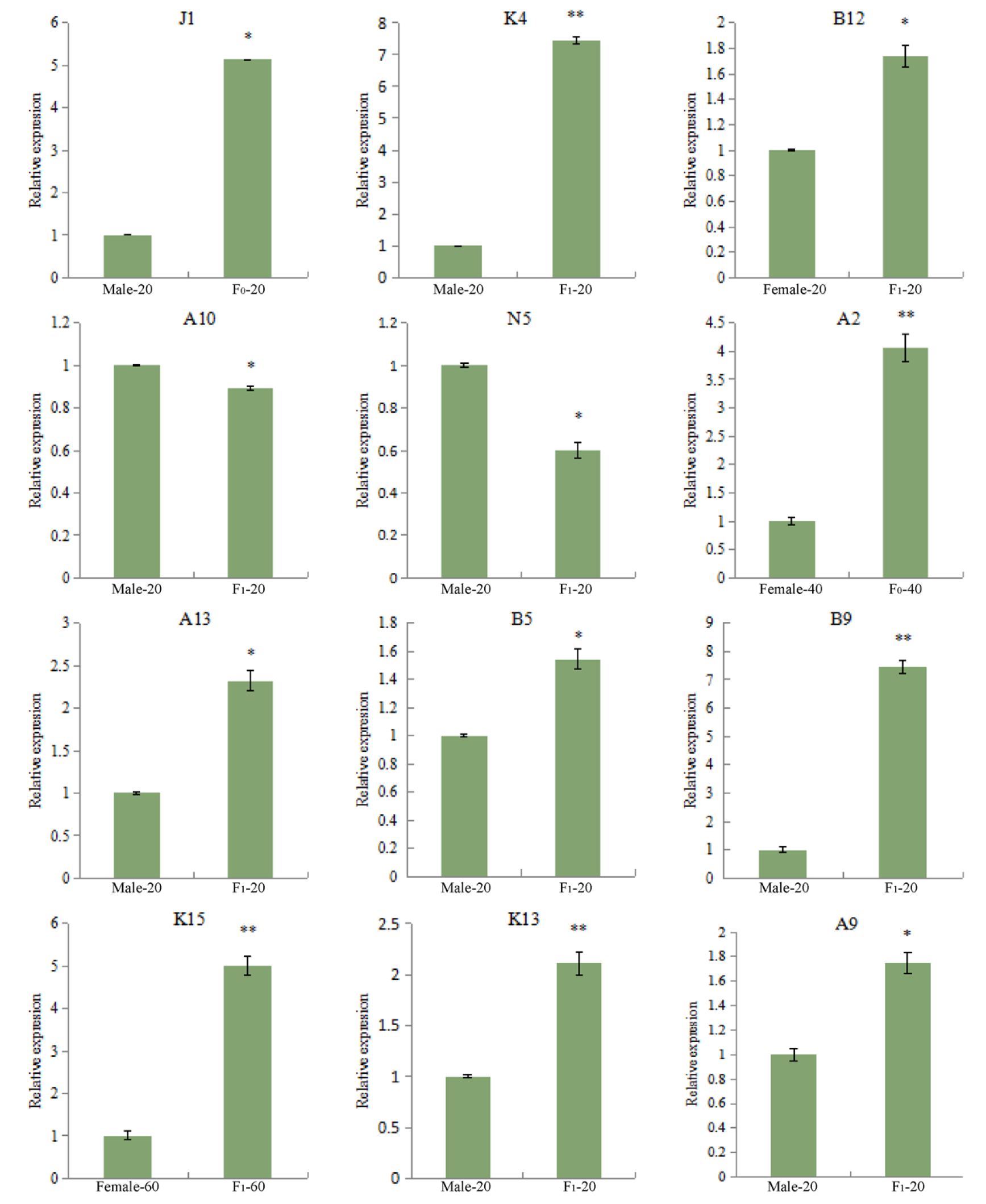
The RT-qPCR Results of Partially Differentially Expressed Proteins between Castor Parents and their Offspring. Note: *: 0.01<P≤0.05, **: 0<P≤0.01.

**Fig. 8-1.**
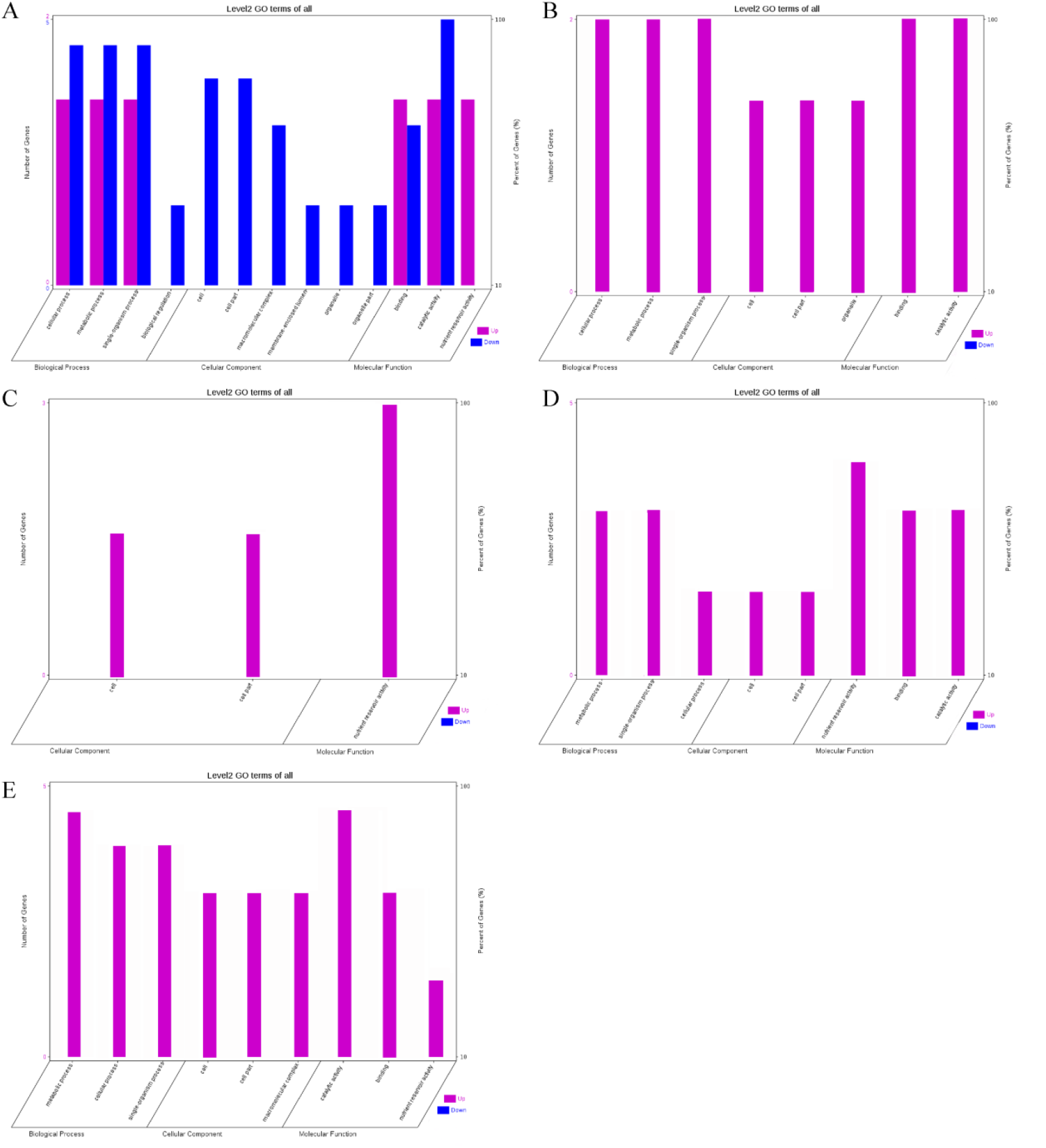
GO Ffunctional Aannotation of Ddifferentially Eexpressed Pproteins in Sseeds at 60 DPP between Ccastor Pparents and their Ooffspring. Note: A: Male, F1, 60 DPP; B: Female, F1, 60 DPP; C: Male, F0, 60 DPP; D: Female, F0, 60d; E: F1 vs. F0, 60 DPP.

### RT-qPCR results of the DEPs

Twelve DEPs in the seeds between castor parents and their offspring (Table 3) were selected for RT-qPCR analysis, and the results are shown in Fig 9. The expression of the genes corresponding to the 12 DEPs all showed differences consistent with those of the DEPs.

**Table 3.** The 12 Differentially Expressed Proteins Chosen for RT-qPCR Analysis.

| Spot No. | Protein Name | Source | Time (d) | Related Traits |
| --- | --- | --- | --- | --- |
| J1 | Stearoyl ACP desaturase | F<sub>0</sub> Male | 20 | Oil synthesis<br>Amino acid metabolism |
| K4 | Oil body associated protein 2C | F<sub>1</sub> Male | 20 | Oil synthesis<br>Amino acid metabolism |
| B12 | Acetyl-CoA carboxylase | F<sub>1</sub> Female | 20 | Oil synthesis<br>Amino acid metabolism |
| A10 | 26S protease regulatory subunit 6a | F<sub>1</sub> Male | 20 | Pollen abortion |
| N5 | Cysteine protease | F<sub>1</sub> Male | 20 | Pollen abortion |
| A2 | Superoxide dismutase | F<sub>0</sub> Female | 40 | Yield and stress resistance |
| A13 | Amylosynthetase | F<sub>1</sub> Male | 20 | Yield and stress resistance |
| B5 | Abundant protein D-7 in late embryonic development | F<sub>1</sub> Male | 20 | Yield and stress resistance |
| B9 | Glutathione peroxidase | F<sub>1</sub> Male | 20 | Yield and stress resistance |
| K15 | Alcohol dehydrogenase 1 | F<sub>1</sub> Female | 60 | Yield and stress resistance |
| K13 | 2S albumin precursor | F<sub>1</sub> Male | 20 | Yield and stress resistance |
| A9 | Aspartic protease | F<sub>1</sub> Male | 20 | Yield and stress resistance |

## Discussion

Through comparative proteomic analysis of seeds at different developmental stages between castor parents and their offspring, 122 DEP spots were detected by MS, though they only contained 93 DEPs. This is because a number of proteins were significantly differentially expressed at all developmental stages of seeds, such as starch synthase, D-7 LEA protein, SOD, and APs.

Most of the DEPs were associated with stress responses (61.29%) and they were mostly upregulated in the seeds of offspring, which was consistent with the results of physiological and biochemical measurements. SOD is an antioxidant enzyme in the reactive oxygen species scavenging system that effectively prevents peroxides (e.g., H_2_O_2_) from damaging the structure and function of plant cells. This enzyme participates in the rcu04146 pathway (peroxisome). The peroxisome, an essential organelle of plants, plays a key role in redox signal transduction and lipid homeostasis [11-15]. APs are mainly involved in such processes as metabolism, precursor protein processing, protein degradation, and programmed cell death, as well as pollen germination and pollen-tube growth in plants [16-18]. D-7 LEA is a stress-responding protein that massively accumulates in the late embryonic stage and is generated under adverse conditions such as drought. It regulates the metabolic balance of plants through mechanisms such as stabilizing the cell membrane and binding metal ions [19-21]. Glutathione peroxidase contains sulfhydryl groups and can effectively block the free radicals of reactive oxygen species from further damaging the body by scavenging H_2_O_2_ and lipid peroxides [22,23]. ADH1, a medium-chain zinc-containing enzyme widely found in various tissues in plants, binds zinc ions. Zinc plays an essential role in the synthesis of indole-3-acetic acid (IAA), an auxin in plants. Zinc deficiency can lead to decreased tryptophan and IAA content, hindering the growth of plant roots. Therefore, ADHs are is closely related to plant growth and development [24-26]. In addition to those five proteins, several DEPs between offspring and parents were associated with stress responses, including 14-3-3-like protein, BADH1 (chloroplast), and l-ascorbate peroxidase.

Among the DEPs between castor offspring and parents, 17.20% were involved in seed nutrient storage, most of which were upregulated in the offspring seeds. Starch synthase is a key enzyme involved in starch synthesis and metabolism. Starch, the main storage carbohydrate, may account for 50% of the dry matter in storage organs (tubers or seeds), and it is an important energy source for plant growth and development, so its content directly determines the yield of crops [27-29]. 2S albumin precursor, a storage protein, can gradually be degraded to produce various amino acids and thereby provide essential nitrogen, carbon, and sulfur sources for plant growth during the seed germination and early growth stages, which is of great significance for plant growth and development [30]. Vicilin GC72-A belongs to the family of antimicrobial peptides, a class of small-molecule polypeptides synthesized by plants to defend against pathogen invasion; these antimicrobial peptides can inhibit or kill a variety of microorganisms and fungi [31,32]. Other DEPs associated with seed nutrient storage of plants included adenosyl homocysteinase, NOP56, and salicylic acid-binding protein 2.

Among the DEPs between castor offspring and parents, 12.90% was related to plant fatty acid synthesis and metabolism, most of which were upregulated in the offspring seeds. This result was consistent with the results of trait and quality assessment. Stearoyl-ACP desaturase is a key enzyme in the conversion between stearic acid and unsaturated fatty acids, so its upregulation may increase the unsaturated fatty acid content in plants, thereby improving their resistance to low temperature [33]. OBAP2C plays an essential role in the seed development of plants. The oil body, a lipid-storing subcellular organellar particle in plant seeds, provides nutrients to plants and is critical for plant growth and development [34-36]. Acetyl-CoA carboxylase is a biotinidase catalyzing the carboxylation of acetyl-CoA to form malonyl-CoA in the initial step of fatty acid synthesis. Fatty acids re membrane lipid components of all cells and an important energy reserve of multicellular organisms. Hence, acetyl-CoA carboxylase, the catalyzer of the first step of fatty acid biosynthesis, plays a critical role in plant growth and development [37-39]. Other DEPs associated with fatty acid synthesis and metabolism included EAR precursor, plastid 3-keto-acyl-ACP synthase I, and hydroxyl-ACP dehydrase.

Only 6.45% of the DEPs between castor offspring and parents were related to amino acid metabolism, most of which were upregulated in the offspring seeds. Both transaldolase and transketolase (chloroplast) are enzymes involved in the nonoxidative phase of the pentose phosphate pathway. Transaldolase catalyzes the transfer of the dihydroxyacetone moiety from sedoheptulose-7-phosphate to glyceraldehyde-3-phosphate; as a result, it can regulate the production of ribose-5-phosphate and NADPH by the pentose phosphate pathway, through which it links the pentose phosphate shunt and glycolysis pathways and regulates the balance between the oxidative and nonoxidative phases of the pentose phosphate pathway. Transketolase is a “nonregulatory” enzyme that catalyzes the reversible transfer of aldehyde group from activated ketose (donor) to activated aldose (receiver). The upregulation of transketolase expression facilitates carbon fixation in plants, as well as plant growth and development [40-44]. SAMS is the only enzyme synthesizing SAM in organisms. SAM, a metabolite from the interaction of methionine and adenosine triphosphate, has been widely found in animals, plants, and microorganisms. It is involved in up to 40 types of biochemical reactions and plays a significant role in improving the drought resistance of plants [45-48]. All three enzymes mentioned above were involved in rcu01230 (biosynthesis of amino acids). In addition, NADP-dependent isocitrate dehydrogenase, l,l-diaminopimelate aminotransferase (chloroplast), and 5-methyltetrahydroproylglutamate-homocysteine methyltransferase in the amino acid biosynthesis pathway were also differentially expressed in seeds between castor offspring and parents.

The high expression of cysteine protease and 26S protease regulatory subunit 6a can lead to abnormal degradation of some regulatory proteins in anther tissue and cause pollen abortion. In this study, these two proteins were downregulated in F_1_ compared with their male parent [49]. Importantly, we found that cysteine protease inhibitor was upregulated in the offspring seeds compared with their parents. This enzyme can bind to cysteine protease to inhibit its activity and reduce the risk of pollen abortion, and it is involved in the stress responses of plants under adverse conditions and participates in physiological processes such as senescence of plants [50].

## Conclusions

To conduct a proteomic analysis to compare the differences in castor seeds at different developmental stages between parents and their offspring, we first optimized our two-dimensional electrophoresis system. The protein atlas obtained from SDS-PAGE following IEF was the best when using samples after impurity removal, 24-cm dry gel strips (pH 3-10 NL), and a protein loading amount of 800 μg.

By 2D-DIGE analysis under these optimal two-dimensional electrophoresis, 93 DEPs were identified in the seeds at different developmental stages between castor parents and offspring. They belonged to five categories: proteins related to stress response (61.29%), seed nutrient storage (17.20%), fatty acid synthesis and metabolism (12.90%), amino acid metabolism (6.45%), and others (2.15%).

RT-qPCR was performed on 12 DEPs, including 1) stearoyl-ACP desaturase, OBAP2C, and acetyl-CoA carboxylase, which were upregulated in the offspring seeds compared with their parents and are involved in oil synthesis and fatty acid synthesis and metabolism; 2) 26S protease regulatory subunit 6a and cysteine protease, which were downregulated in the offspring seeds compared with their parents and are involved in pollen abortion; and 3) SOD, starch synthase, D-7 LEA protein, glutathione peroxidase, ADH1, 2S albumin precursor, and APs, which were upregulated in the offspring seeds compared with their parents and are involved in plant resistance to adverse conditions. The RT-qPCR results showed that the expression levels of the genes encoding these 12 DEPs were consistent with the expression of the DEPs.

In summary, this study can help guide the production of castor beans and lays the foundation for cultivating new castor varieties with high yield, high oil content, and strong stress resistance.

